## Supplementary Figure 1 for "Rapid, DNA-induced interface swapping by DNA gyrase"

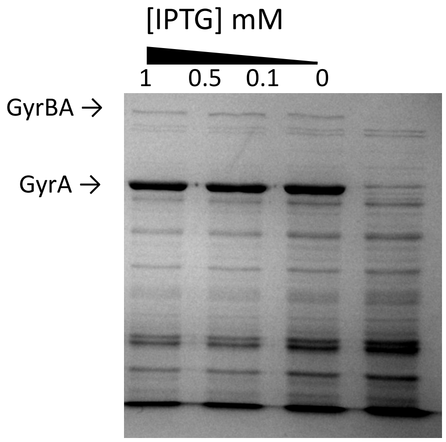


**a**

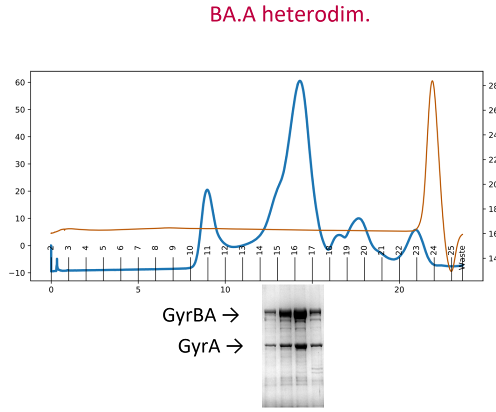


**b**

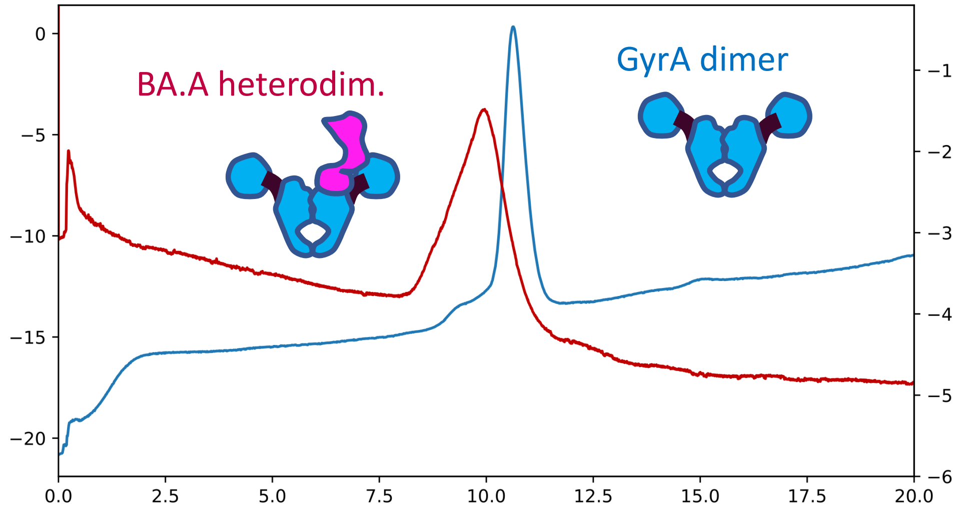


**c**

**Supplementary Figure 1.** Gel filtration and SDS-PAGE analysis of our heterodimer preparation. **a.** Co-expression of GyrBA fusion and GyrA in *E. coli*. GyrA is expressed at a much higher level compared to GyrBA, thereby favouring the formation of BA.A heterodimers and $A_2$ dimers. **b.** Last polishing step of the heterodimer preparation; the preparation is passed through a Superose 6 column. The abscissa is the elution volume. In blue is the UV absorbance in arbitrary units; in brown is the conductivity (the salt peak indicates the total column volume). The fractions are indicated at the bottom. Fractions 10 to 20 contain the heterodimer complex and are analyzed by SDS-PAGE (bottom) stained with Coomassie. The GyrA and GyrBA fusion bands are indicated. **c.** Analytical gel filtration of BA.A (red line) compared to a GyrA sample (blue line); an analytical Superdex 200 column was used, BA.A and GyrA form distinct peaks. The GyrA peak elutes at around 160 kDa (compared to a native marker) and the BA.A heterodimer at around 180-190 kDa, much lower than its theoretical MW.
