## Supplementary Figure 2 for "Rapid, DNA-induced interface swapping by DNA gyrase"

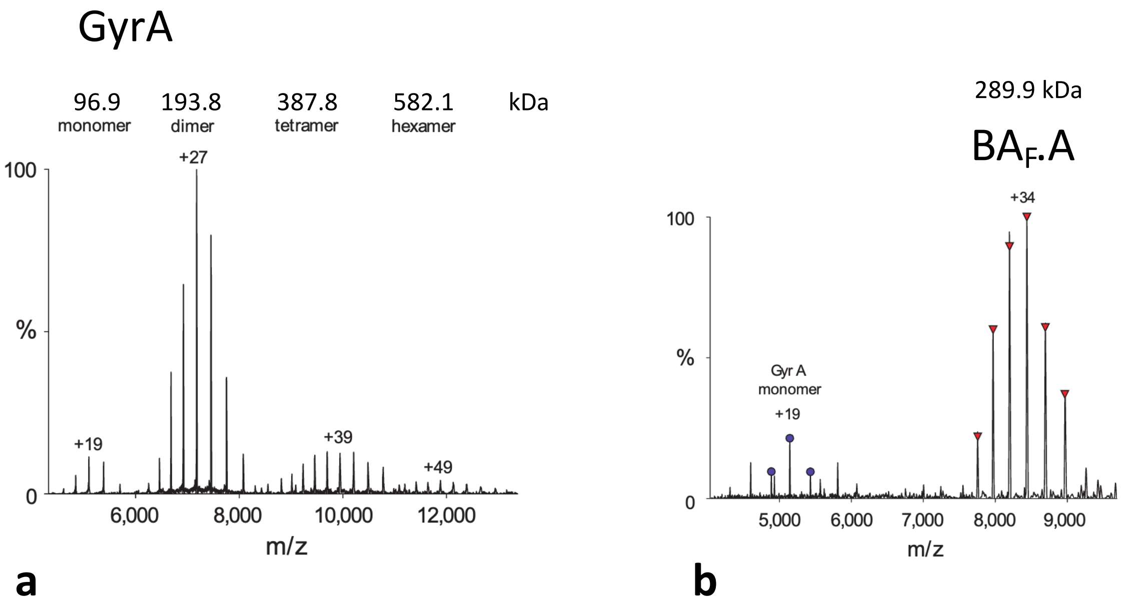


**Supplementary Figure 2.** Analysis of GyrA and $BA_F$.A preparation by native mass spectrometry. **a.** GyrA. The preparation resolves into four peaks with Molecular Weights (MWs) consistent for the GyrA monomer, dimer, tetramer and hexamer. Measured MWs are indicated in kDa. The theoretical MW of the GyrA monomer is 96,947 Da. **b.** $BA_F$.A. The preparation resolves into two peaks, with some impurities at around 60 kDa. The measured MW of $BA_F$.A is indicated in kDa. The theoretical MW of $BA_F$.A is 285,847 Da. The numbers indicate the ionization state of the most intense peak.
