## Supplementary Figure 3 for "Rapid, DNA-induced interface swapping by DNA gyrase"

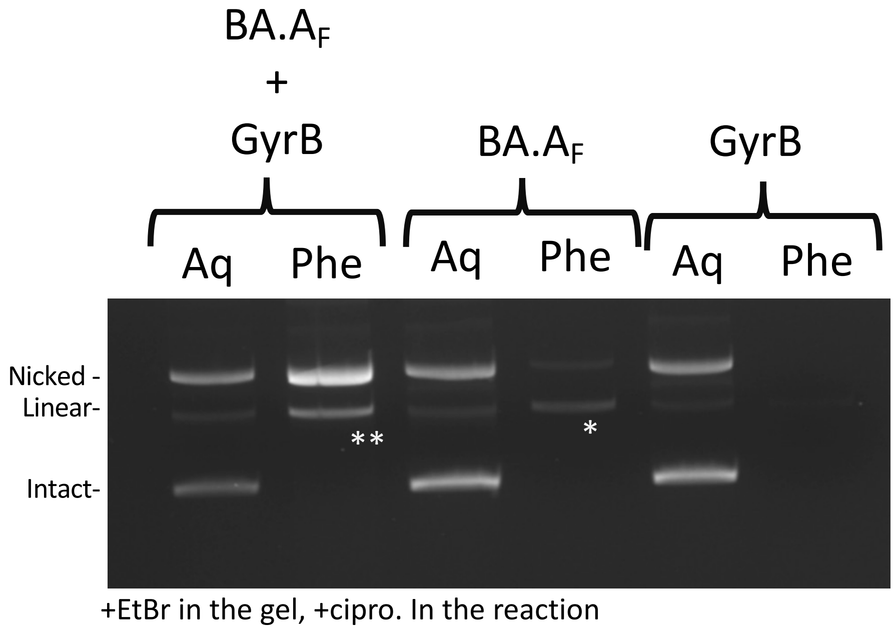


**Supplementary Figure 3.** BA.A$_F$ can form double-strand cleavage complexes in the presence of GyrB. 5 pmole of BA.A$_F$ were incubated in a cleavage assay in the presence of 8 pmole of GyrB. Control cleavage assays with BA.A$_F$ and GyrB alone were also performed. Cleavage complexes were purified and separated from naked DNA by phenol extraction (Supp. Methods). Protein-DNA adducts are trapped at the phenol interface and can be recovered (Phe). Naked DNAs remain in the aqueous phase (Aq). In the presence of GyrB, BA.A$_F$ produces mostly single-strand cleavage complexes, as expected. However, a significant minority of double-strand cleavage complexes are also recovered. The ** band is significantly more intense than the * band (as ascertained by densitometry) constituted by background double-strand cleavage complexes that occurs with BA.A$_F$ alone (probably from contaminating GyrBA fusion dimers). The origin of these GyrB-induced double-strand cleavage complexes is discussed in the main manuscript. This experiment was reproduced three times with virtually identical results.
