## Supplementary Figure 4 for "Rapid, DNA-induced interface swapping by DNA gyrase"

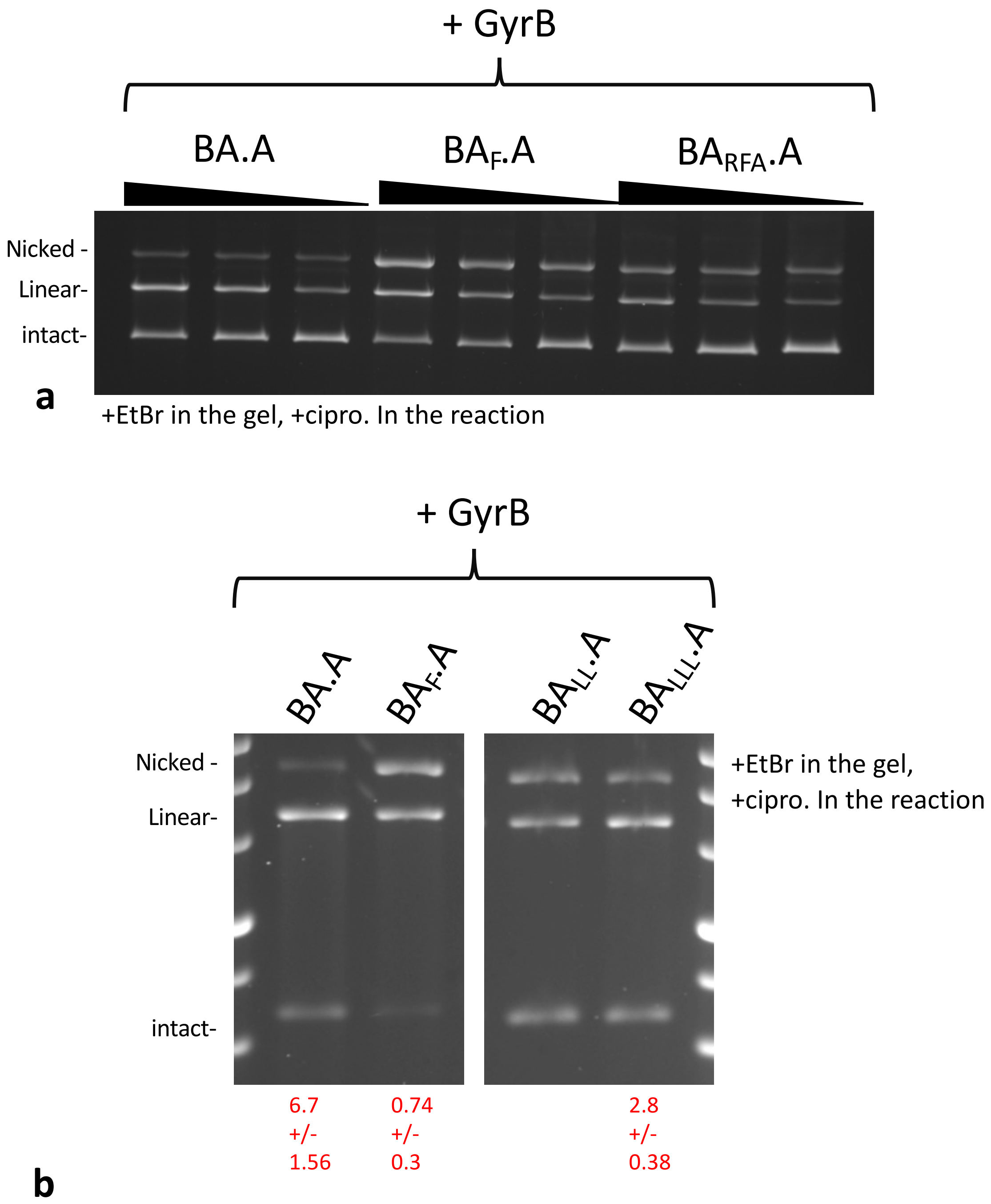


**Supplementary Figure 4.** Cleavage activity of various heterodimers mutant. **a.** 5, 2.5 and 1.25 pmole (triangle indicate increasing dose) of BA.A, $BA_{F}$.A and $BA_{FA}$.A (where the tyrosine adjacent to the catalytic tyrosine is mutated to an alanine) were incubated in a cleavage assay in the presence of GyrB. $BA_{FA}$.A displays significantly more double-strand to single-strand cleavage. **b.** 5 pmole of $BA_{F}$.A, $BA_{LL}$.A and $BA_{LLL}$.A were tested as above. Again $BA_{LL}$ .A and $BA_{LLL}$.A displayed higher double-strand cleavage activity (this is especially marked for $BA_{LLL}$.A). The double- to single-strand cleavage ratio is indicated in red with the standard deviation (unbiased) from three independent experiments.
