## Supplementary Figure 5 for "Rapid, DNA-induced interface swapping by DNA gyrase"

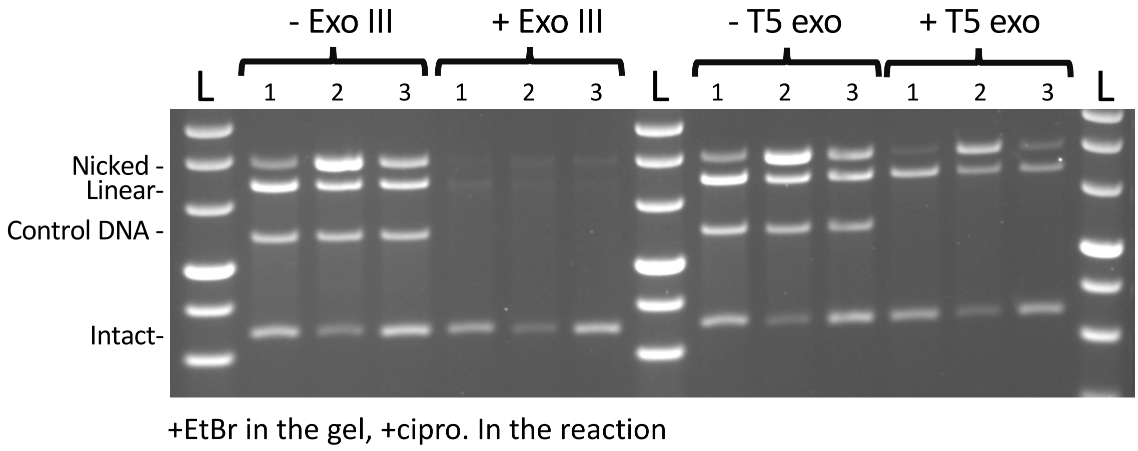


**Supplementary Figure 5.** Exonuclease sensitivity of heterodimeric gyrase cleavage products. All reactions were performed with 5 pmole of heterodimer and 8 pmole of GyrB and purified before treatment (or not) with exonuclease as indicated. A purified PCR fragment was added to all reactions to control for nuclease activity (Control DNA). 1: BA.A (wild-type). 2: $BA_F$.A. 3: $BA_{LLL}$.A. L: Ladder
