## Supplementary Figure 6 for "Rapid, DNA-induced interface swapping by DNA gyrase"

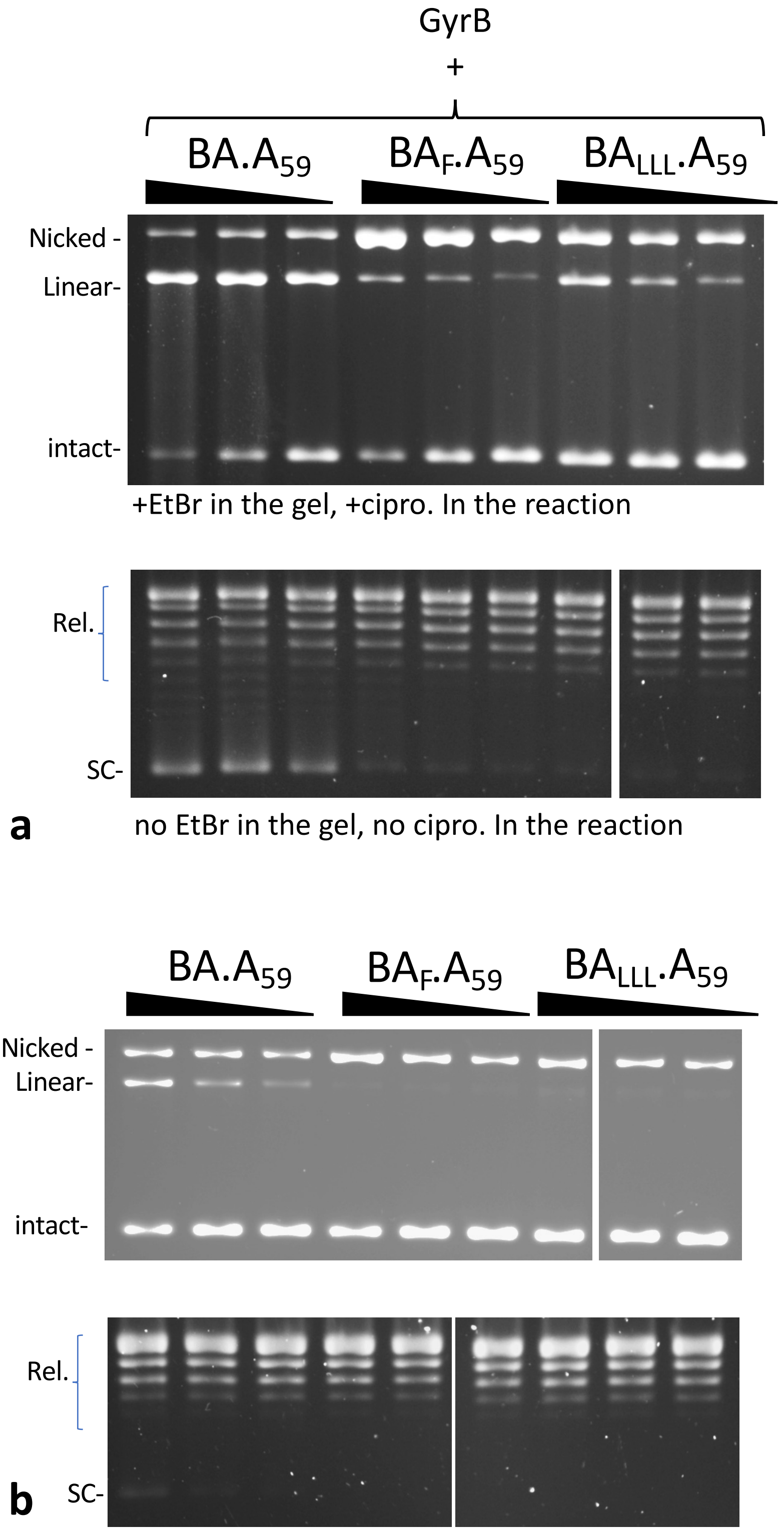


**Supplementary Figure 6. a.** Cleavage and supercoiling activity of various heterodimer mutants, lacking a CTD. **top.** 5, 2.5 and 1.25 pmole (triangle indicate increasing dose) of $BA.A_{59}$, $BA_F.A_{59}$ and $BA_{LLL}.A_{59}$ were incubated in a cleavage assay in the presence of GyrB. $BA_{LLL}.A_{59}$ displays significantly more double-strand to single-strand cleavage compared to $BA_F.A_{59}$. **bottom.** 1.25, 0.625 and 0.312 pmole pmole of $BA.A_{59}$, $BA_F.A_{59}$ and $BA_{LLL}.A_{59}$ were tested in a supercoiling assay in the presence of GyrB. **b. top.** Cleavage assay, as above, in the absence of GyrB. **bottom.** Supercoiling assay, as above, in the absence of GyrB.
