## Supplementary Figure 7 for "Rapid, DNA-induced interface swapping by DNA gyrase"

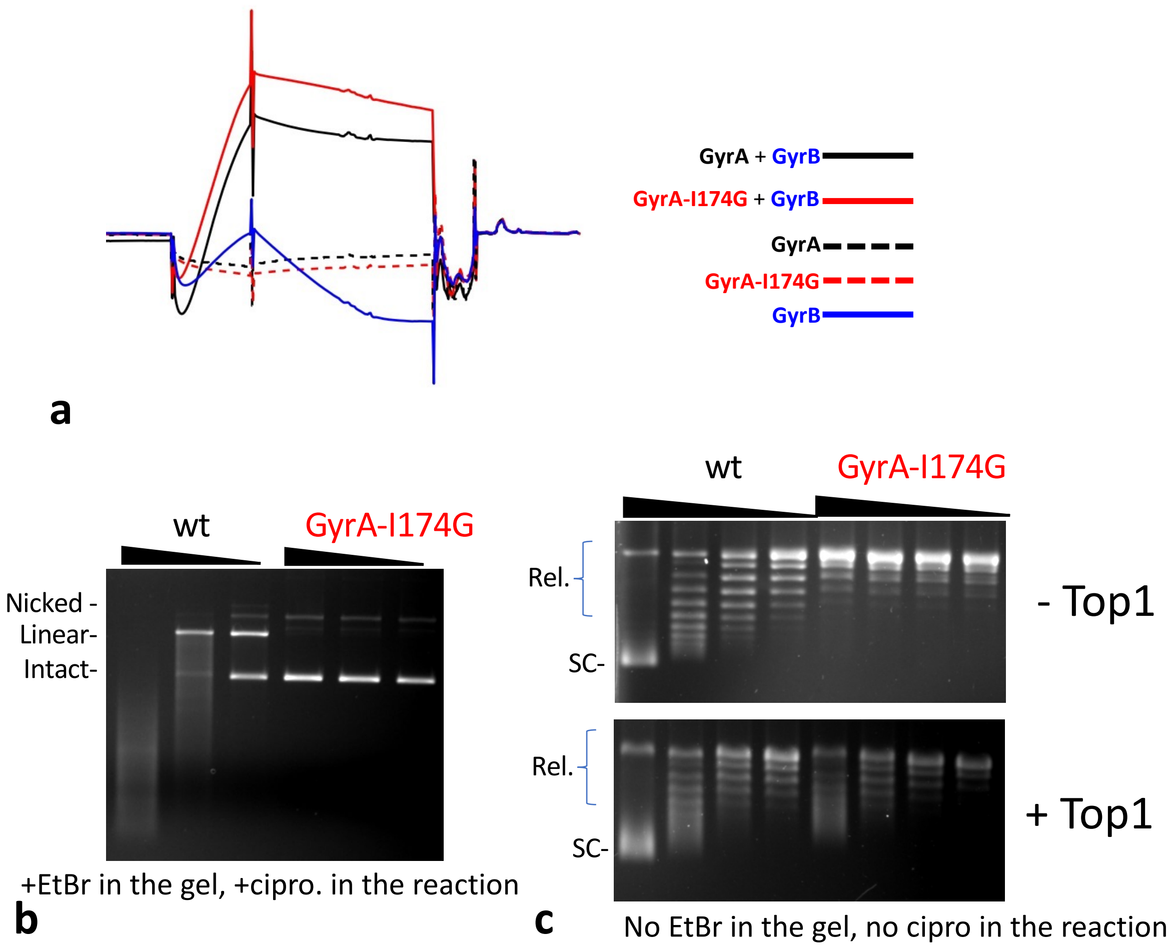


**Supplementary Figure 7.** Analysis of cleavage and DNA binding properties of the GyrA-I174G mutant homodimer. **a.** Surface Plasmon Resonance performed with a 170 bp DNA fragment immobilized on a SPR chip. Various preparations were flowed on to the chip, as indicated and the response recorded. GyrA dimers (wild-type and mutant) showed no binding on their own. GyrB showed limited binding. GyrA and GyrB in combination displayed a strong response, showing robust binding. The concentration of GyrA dimers and GyrB were identical between experiments. **b.** Cleavage activity of wild-type GyrA and GyrA-I174G. Increasing amounts (triangle) of GyrA dimers as indicated were incubated in the presence of GyrB and 20 µM ciprofloxacin. The I174G mutation abolished cleavage. Supercoiling activity is also abolished (not shown). **c.** Topological footprint of GyrA wild type and mutant. Increasing amounts of GyrA (triangle) were incubated, in the absence of ATP and ciprofloxacin, with relaxed plasmid DNA and GyrB. *E. coli* Top1 was then added to relax free supercoiling, deproteinization reveals the positive supercoiling constrained by the enzyme. Wild type gyrase can relax DNA, and therefore constrained positive supercoils are visible in the absence of Top1. GyrA-I174G has lost cleavage activity and therefore relaxation activity but can still constraint a limited number of positive supercoils. The handedness of supercoils was ascertained by running agarose gels in the presence of chloroquine (not shown).
