## Supplementary Figure 8 for "Rapid, DNA-induced interface swapping by DNA gyrase"

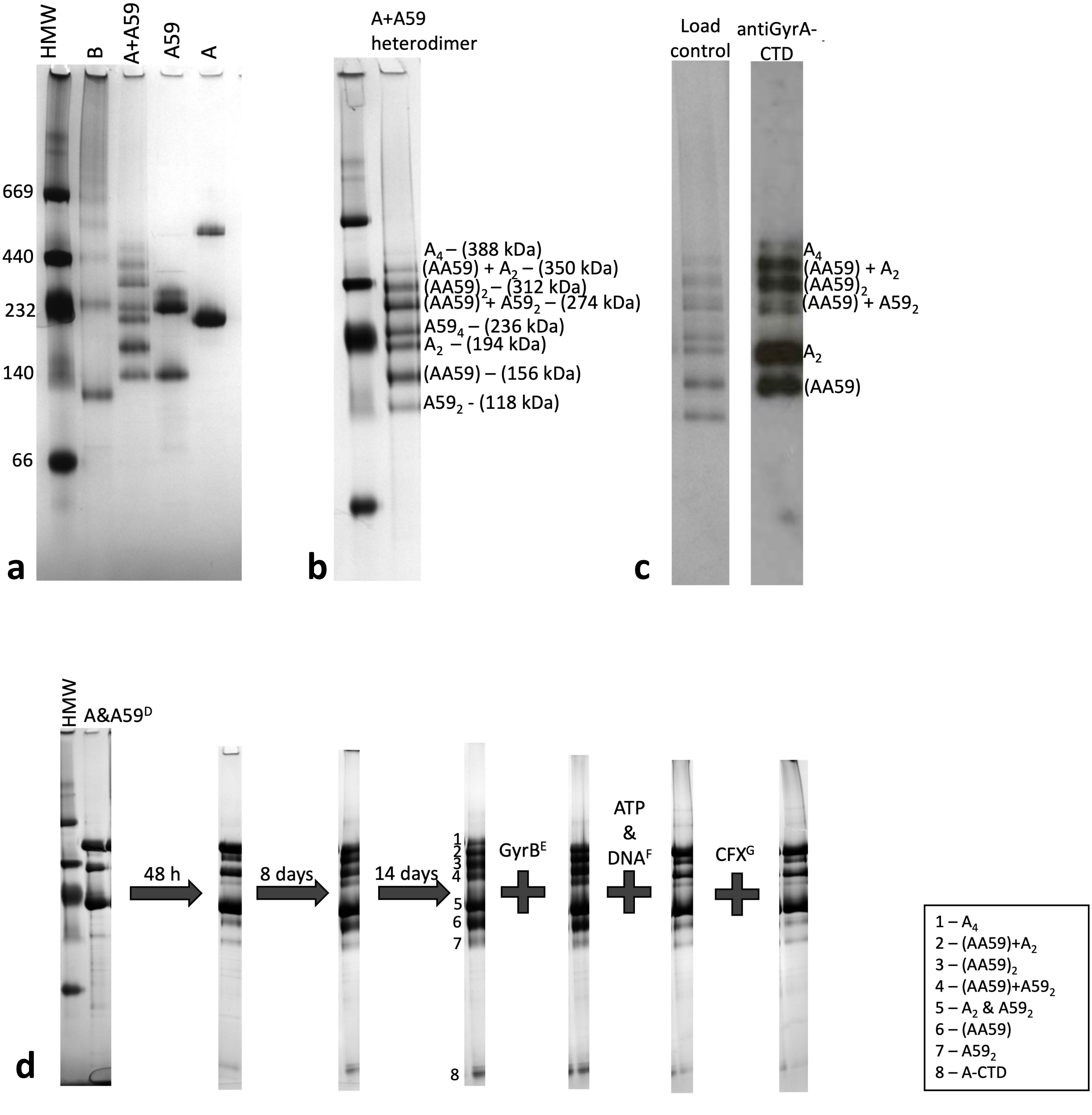


**Supplementary Figure 8.** Blue-Native PAGE and Western blot of the GyrA/GyrA59 heterodimers. **a.** BN-PAGE GyrA (A), GyrA$_{59}$ (A59), a refolded heterodimer (A+A59) and GyrB (B) run on a 4-12% gradient gel. HMW is the high molecular weight marker with the size of each band in kDa down the left-hand side. **b.** Refolded heterodimer run as in **a** with the higher-order complexes highlighted alongside, with their predicted molecular weights. **c.** Refolded heterodimer run as in **a** stained with coomassie alongside the western blot probing for the GyrA-CTD. This antibody binds to full length GyrA but not A$_{59}$. The bands were assigned on the basis of their reactivity with the antibody, combined with their molecular weight, estimated by comparison to the marker. A$_4$ is the GyrA tetramer, A59$_4$ is the GyrA59 tetramer, (AA59)$_2$ is the GyrA/GyrA59 heterotetramer, (AA59) is the heterodimer, A$_2$ is the GyrA dimer, A59$_2$ is the GyA59 dimer. **d.** Effect of long-term incubation on subunit exchange. The A&A59 sample is run as in **a** after incubation for the indicated length of time, up to 14 days. 1 – 8 indicate the GyrA/GyrA59 complexes species. The arrow indicates the position of the heterodimer A.A59 (species 6). The last time point of the same experiment performed with the addition of GyrB, GyrB + ATP +DNA and GyrB + ATP +DNA + ciprofloxacin is shown on the right.
