## Supplementary Figure 9 for "Rapid, DNA-induced interface swapping by DNA gyrase"

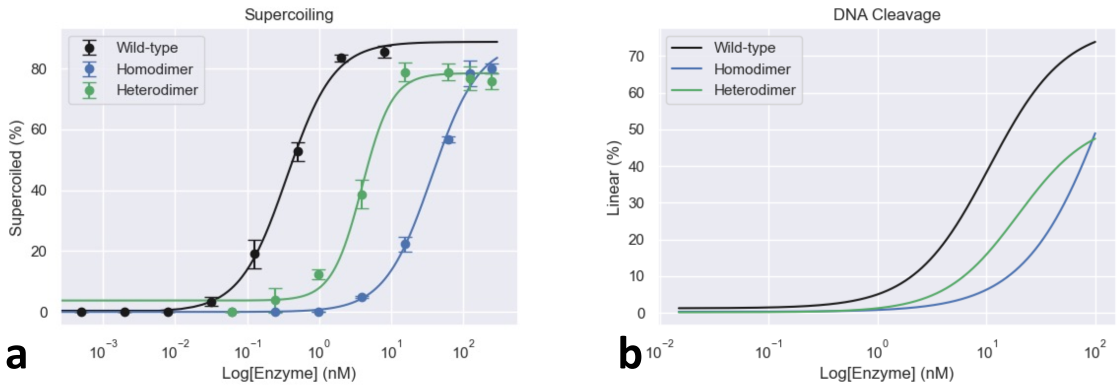


**Supplementary Figure 9.** Comparison of the activity of the GyrA dimer + free GyrB (wild type), BA.A heterodimer + free GyrB (heterodimer) and $(BA)_2$ (homodimer). **a.** Supercoiling assay. The $IC_{50}$ (concentration of enzyme producing half the observed supercoiling) is 0.36 nM for the GyrA dimer (wild type), 4 nM for the heterodimer and 36.7 nM for the $(BA)_2$ homodimer. The cleavage did not plateau for the heterodimer and $(BA)_2$ homodimer, precluding $CC_{50}$ measurements (concentration of enzyme producing half the maximum level of cleavage).
