## Supplementary Figure 10 for "Rapid, DNA-induced interface swapping by DNA gyrase"

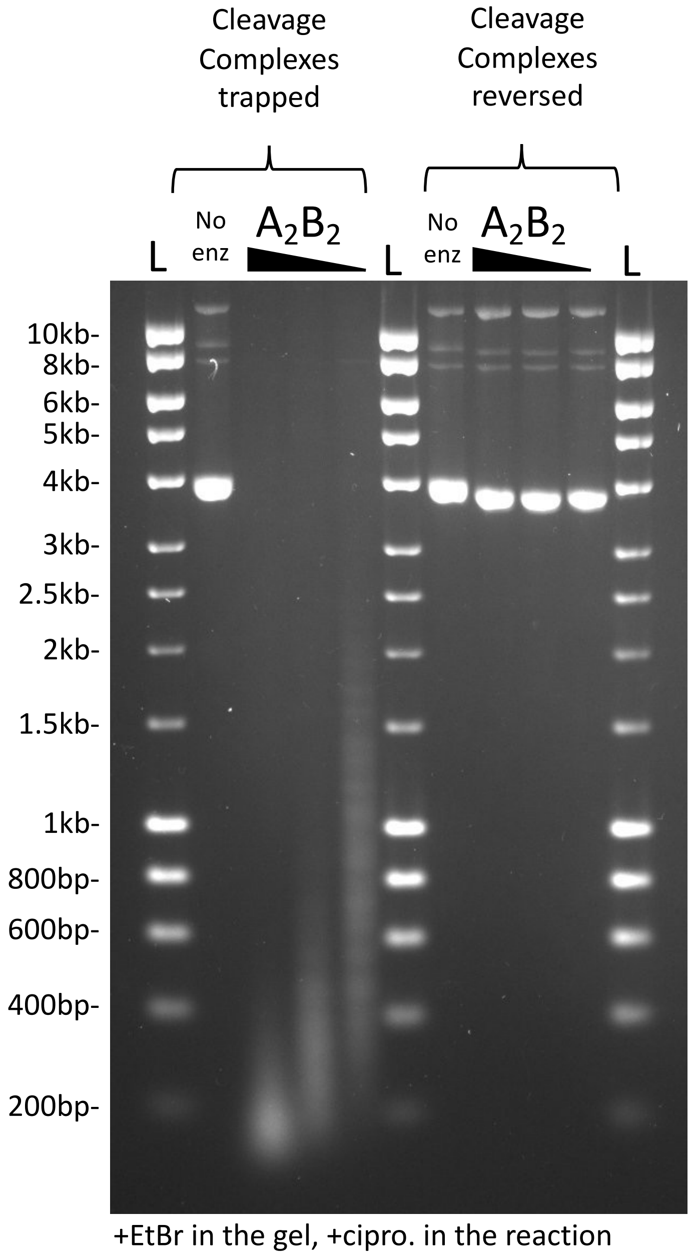


**Supplementary Figure 10. a.** Gyrase can stabilize cleavage complexes ~200 bp apart on DNA. A cleavage assay was performed with either no enzyme or 30, 15 and 7.5 pmole of the $A_2B_2$ tetramer on a ~8kb plasmid (pIRT2tg-NAT), in the presence of 20 µM ciprofloxacin. Cleavages complexes were either trapped by the addition of SDS (methods) or reversed by the addition of EDTA, followed by Proteinase K digestion. The observed cleavage is entirely reversible, showing it arises from the entrapment of genuine cleavage complexes. L: DNA ladder.
