## Supplementary Figure 11 for "Rapid, DNA-induced interface swapping by DNA gyrase"

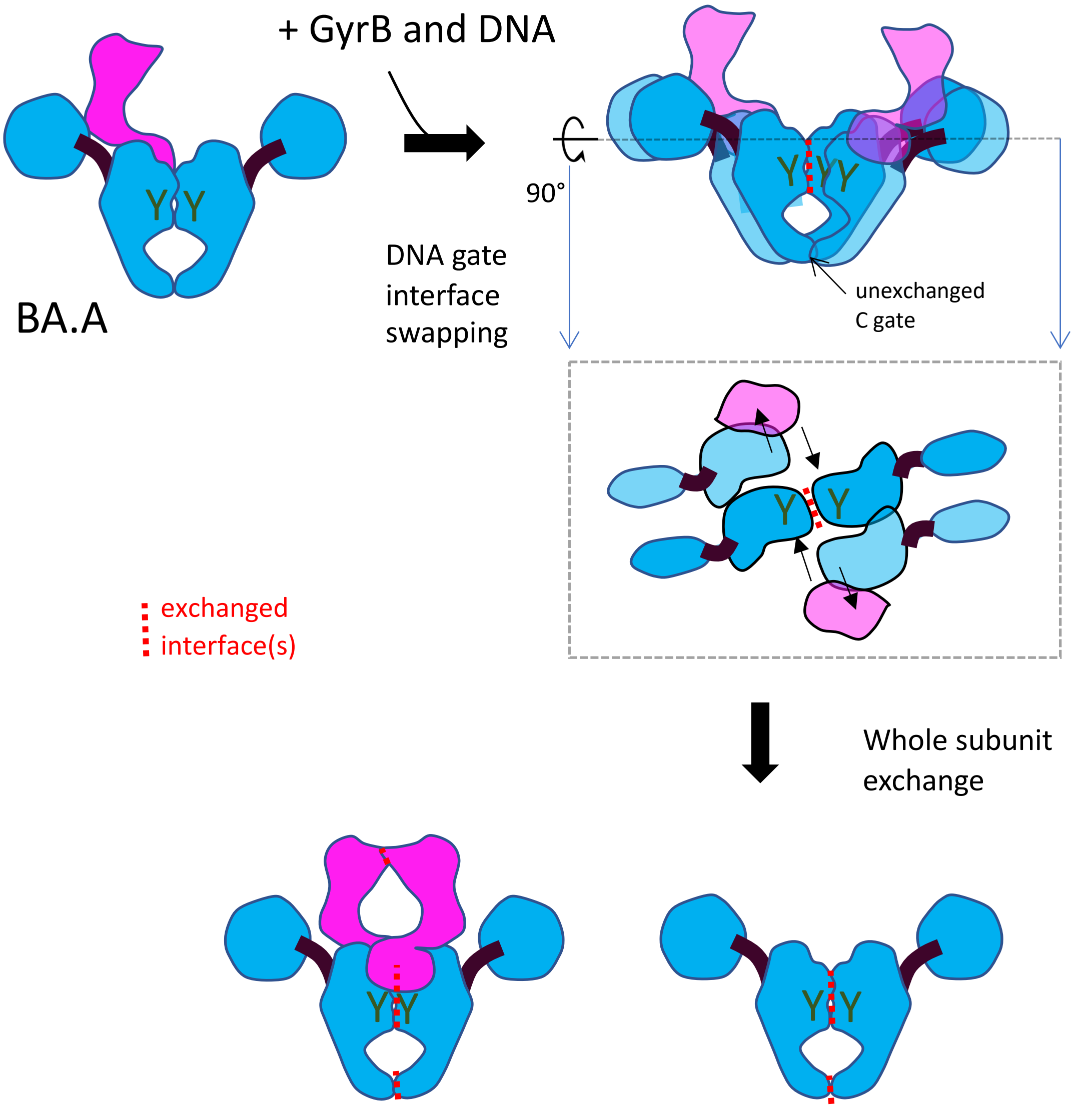


**Supplementary Figure 11.** Schematic of possible sequential interface swapping mechanism leading to complete subunit exchange. Top left, schematic of the BA.A heterodimer. Two heterodimers having undergone DNA gate interface swapping are represented Top Right. The exchanged interfaces are shown with the red dashed line. The dashed black line represent a plane intersecting the swapped heterodimer horizontally. Just below the Top Right schematic is a view from the top of this plane and any elements outside the plane is excluded. Bottom. Subsequent exchange of the C gate and the N gate produces two exchanged dimers ($A_2$ and $BA_2$) from the original 2 BA.A heterodimer. The approximate position of the catalytic tyrosine (Y in dark green) is indicated.
