## Supplementary Figure 13 for "Rapid, DNA-induced interface swapping by DNA gyrase"

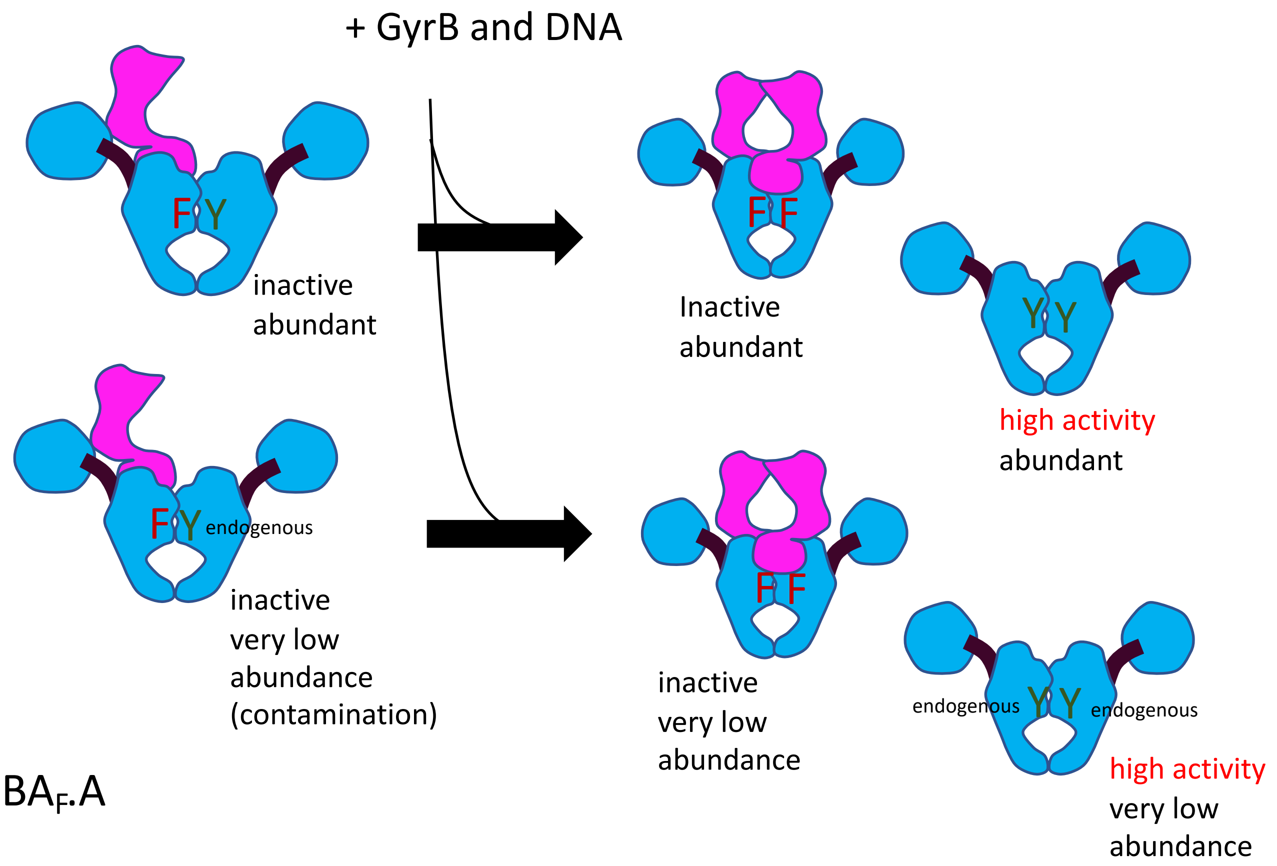


**Supplementary Figure 13.** Schematic of possible sequential interface swapping mechanism leading to complete subunit exchange for the BA$_F$.A preparation. Only complete subunit exchange is shown. The phenylalanine that replaces the catalytic tyrosine is shown in dark red. The estimated activity and abundance of each product is shown. This preparation is therefore expected to have an overall **high supercoiling activity and high cleavage activity.**
