## Supplementary Figure 14 for "Rapid, DNA-induced interface swapping by DNA gyrase"

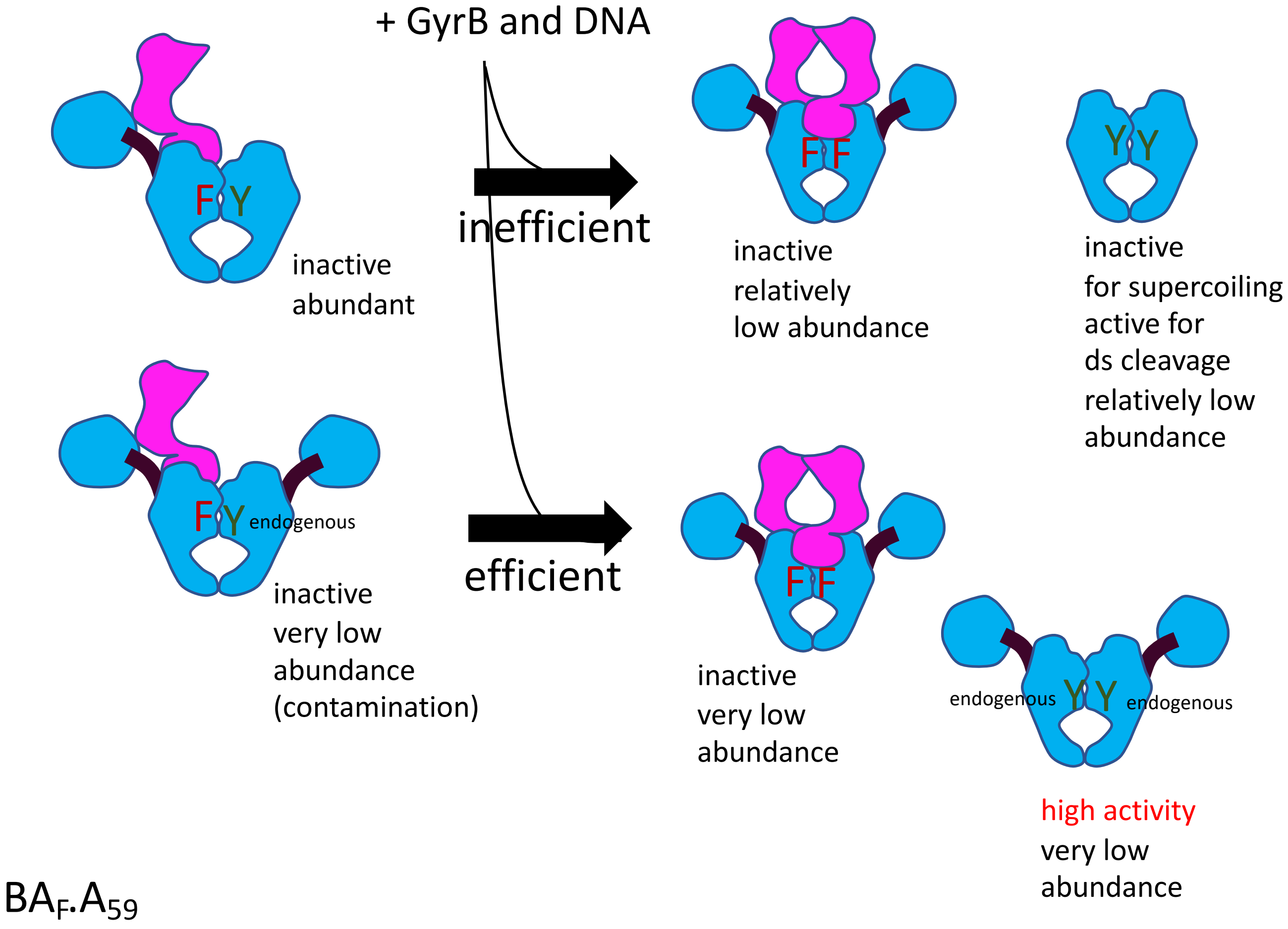


**Supplementary Figure 14.** Schematic of possible sequential interface swapping mechanism leading to complete subunit exchange for the $BA_F.A_{59}$ preparation. Only complete subunit exchange is shown. The phenylalanine that replace the catalytic tyrosine is shown in dark red. The estimated activity and abundance of each product is shown. This preparation is therefore expected to have and overall **low supercoiling activity, low cleavage activity.**
