## Supplementary Figure 15 for "Rapid, DNA-induced interface swapping by DNA gyrase"

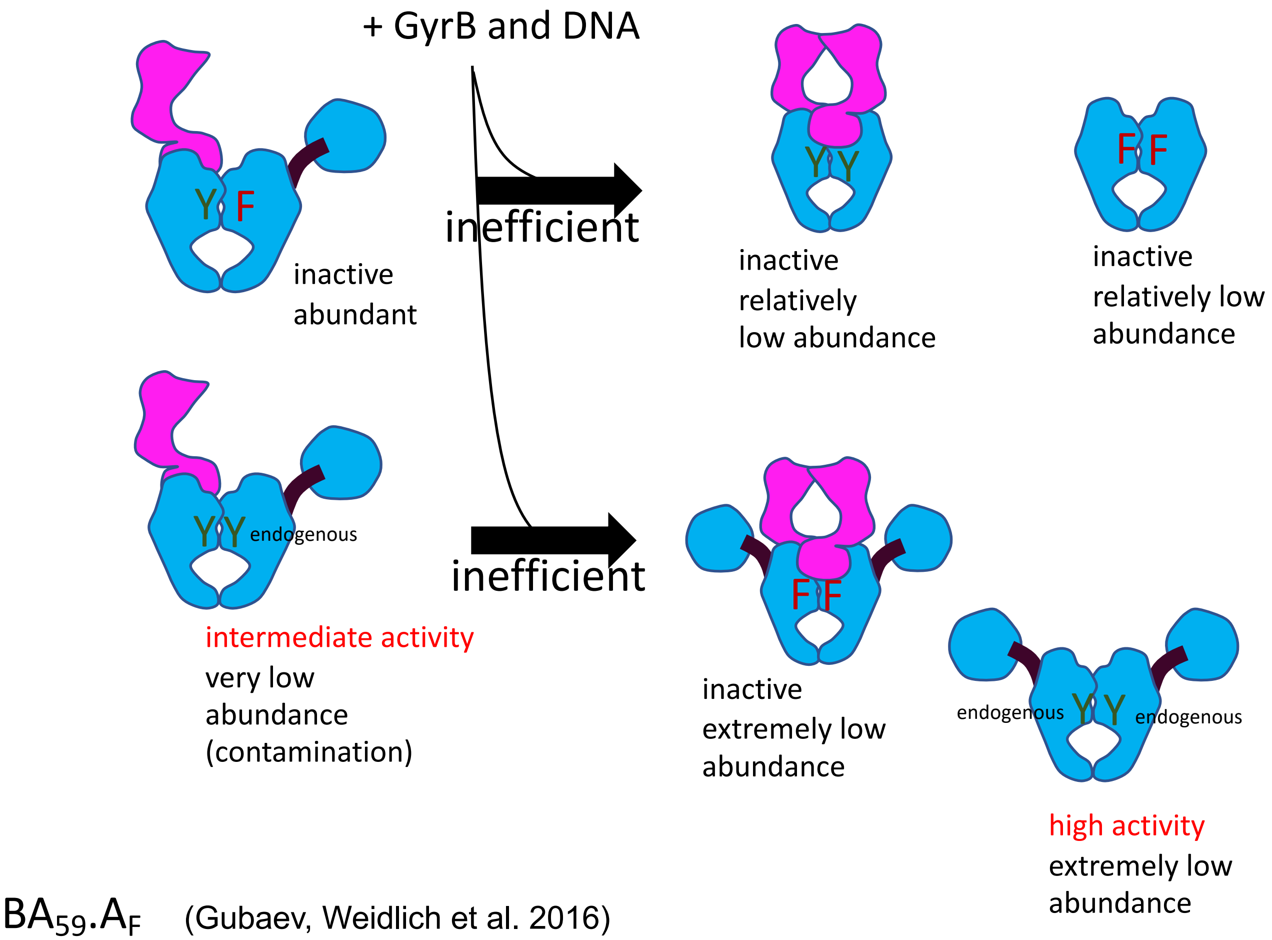


**Supplementary Figure 15.** Schematic of possible sequential interface swapping mechanism leading to complete subunit exchange for the $BA_F.A$ preparation. Only complete subunit exchange is shown. The phenylalanine that replaces the catalytic tyrosine is shown in dark red. The estimated activity and abundance of each product is shown. This preparation is therefore expected to have overall **very low supercoiling activity and low cleavage activity.**
