## Supplementary Figure 16 for "Rapid, DNA-induced interface swapping by DNA gyrase"

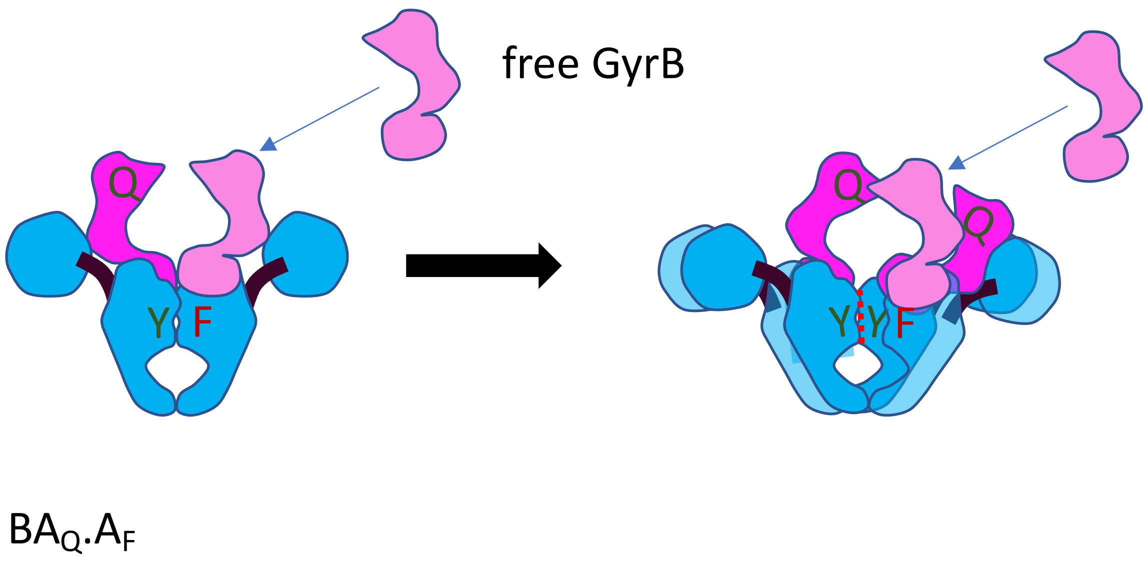


**Supplementary Figure 16.** Schematic of possible DNA gate-only interface swapping mechanism leading to a $BA_Q.A_F$ preparation having some supercoiling activity in the presence of GyrB. The phenylalanine that replace the catalytic tyrosine is shown in dark red. The free GyrB (light pink) can associate with the $GyrA_F$ subunit and dimerise with $GyrB_Q$ from the BA fusion involved in the exchanged DNA gate interface (red dashed line). The ATPase domain of the other $BA_Q$ fusion involved in the exchanged interface. The TOPRIM domain might not even be exchanged and can still promote cleavage. It is also possible that the whole $GyrB_Q$ domain dissociate from the GyrA domain of the fusion, only staying attached to GyrA at the fusion point; the free GyrB could then replace the fused $GyrB_Q$ and interact with the GyrA domain of the fusion normally. The efficiency of this exchange would presumably depend on the flexibility of the fusion point.

In the PowerPoint file this section has a different font
