## Supplementary Figure 17 for "Rapid, DNA-induced interface swapping by DNA gyrase"

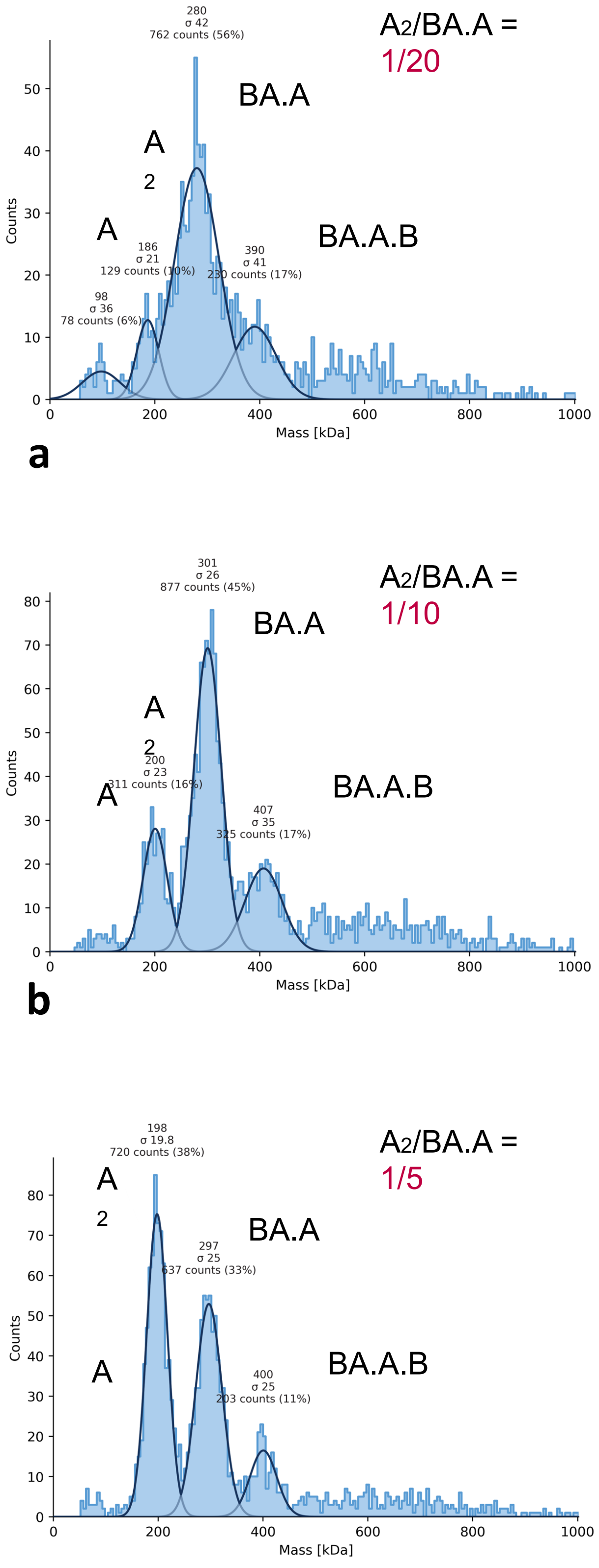


**Supplementary Figure 17.** BA.A heterodimer and GyrA dimer mixing experiment; done to assess sensitivity of mass photometry to contaminating GyrA dimers mixed with BA.A heterodimers. 50 nM of BA.A were mixed with 2.5 nM (**a**), 5 nM (**b**) and 10 nM (**c**) of purified GyrA dimer. Data shows that 1/20 GyrA/BA.A contamination is detectable (**a**). Increasing this ratio drastically increase the count for the GyrA dimer peak to above the BA.A peak (**b** and **c**), despite the latter species being more abundant. This shows that the counts for each species is dependent not only on their abundance, but also how efficiently they collide to the surface. The intensity of peaks should not therefore be used as a measure of abundance.
