## Supplementary Discussion for "Rapid, DNA-induced interface swapping by DNA gyrase"

**Low contamination of BA.A_F_ and BA_F_.A_59_ with endogenous GyrA**

The reconstitution of double-strand cleavage correlates with the reconstitution of supercoiling activity.

However, since the supercoiling reaction is extremely efficient, as opposed to cleavage, a very low amount of contaminating GyrA dimers, below detection levels, could explain the detection of low level of negative supercoiling (see Main Discussion). In the case of BA_F_.A the level of supercoiling is very high, comparable to the wild-type, and cannot be explained by a sub-detection level of GyrA. However, In the case of BA.A_F_, a low level of GyrA dimer, encoded by the endogenous *gyrA* gene could explain the low level of supercoiling we observe. In the case of our heterodimer preparations, both GyrA (untagged) and GyrBA fusion (his-tagged) are co-expressed in *E. coli*. Various dimeric combinations are therefore obtained in the crude extract, before purification. Since the nickel column step pulls down only the GyrBA fusion, GyrA dimers are purified out. It is also possible that some heterodimers contain the His-tagged GyrBA fusion dimerized with a GyrA subunit encoded by the endogenous *gyrA*, even though the expression level of recombinant GyrA is very high (Supp. Fig. 1a), which should minimize their formation. Nonetheless, such complexes are expected to occur during expression and to be co-purified. We observed a low level of GyrA cleavage-dependent labeling with our BA.A_F_ preparation, which demonstrate the existence of these complexes, which we denote BA.A_endogenous_. These complexes can explain the low level of supercoiling observed with the BA_F_.A_59_ heterodimer, which cannot reconstitute supercoiling activity by subunit exchange, since the contaminating BA_F_.A_endogenous_ would reconstitute a GyrA_endogenous_ dimer, which is active in the presence of GyrB. It could also underlie the low level of supercoiling observed with the BA.A_F_ heterodimer in the presence of GyrB, although such activity could also arise from the reconstituted BA fusion dimer. In (Gubaev, Weidlich et al. 2016) no experiments were done to address whether cleavage originated from the expected recombinant subunit as opposed to possible minor contamination with unmutated, endogenous subunits.

Our purification system is not as stringent as in (Gubaev, Weidlich et al. 2016). We use a single tag on the BA fusion and the expression strategy minimizes the formation of unwanted heterodimers with endogenous GyrA. However, we account for contamination and have established that they are minimal thereby reconstituting a weak supercoiling activity. In (Gubaev, Weidlich et al. 2016) an extra tag on GyrA is used for purification and should further minimize endogenous contamination. However, a small contamination is not excluded. We suggest a cleavage radiolabeling of subunits could have been done to address this. If the recombinant GyrA is mutated for cleavage no labeling should occur and the observation of even a very minor GyrA signal suggests contamination. Our GyrA preparations are usually not contaminated by endogenous GyrA. For instance, our preparation of GyrA_I174A_ does not show any supercoiling activity (nor cleavage) in the presence of GyrB. Our GyrB preparation on the other hand can be contaminated by a low amount of GyrA, undetectable with monoclonal antibodies, as evidenced by the very low level of supercoiling induced by GyrB alone at high concentration (in assays, lower concentrations are used to prevent this unwanted activity). This amount of contaminant GyrA can vary from preparation to preparation. It is therefore possible that the heterodimer preparation is contaminated by a very low amount of endogenous GyrA that escapes the tandem tag purification, the GyrB domain of the fusion interacting with GyrA. In addition, we have demonstrated that GyrB oligomerizes. Therefore, two heterodimers such as BA_59_.A_F_ and BA_F_.A_endogenous_ could oligomerize through their GyrB domain (which is unmutated) and the tandem tag purification of BA_59_.A_F_ could indeed co-purify a small amount of BA_59_.A_endogenous_. In (Gubaev, Weidlich et al. 2016) the authors trust that the tandem purification procedure will not produce contamination important enough to explain their results without really testing it. Their only experiment directly addressing contamination is described in their Supplementary Figure 6 (Gubaev, Weidlich et al. 2016). In this experiment it is shown that B*.* subtilis GyrB can produce active gyrase with an E*.* coli GyrA dimer. However, E*.* coli GyrB cannot reconstitute an active gyrase with B*.* subtilis GyrA dimer (ref). They then add either B*.* subtilis or E*.* coli GyrB to a B*.* subtilis BA.A_F_ heterotetramer and show that only the B*.* subtilis GyrB is capable of reconstitute supercoiling activity. They conclude, rightfully, that their B*.* subtilis heterodimer preparation, expressed in E*.* coli, is not contaminated by E*.* coli GyrA dimers, since these would reconstitute active gyrase with added E*.* coli GyrB. However, this experiment does not exclude the presence of BA.A_endogeous_ where the BA fusion is B*. subtilis* and the GyrA subunit is endogenous *E.* coli from the expression strain. This complex could still undergo subunit exchange since the BA fusion side is B subtilis, producing an E*.* coli GyrA dimer that can reconstitute gyrase activity in the presence of B*.* subtilis GyrB, as shown in their previous experiment. When E. coli GyrB is added to the B*.* subtilis BA.A_endogenous_ it cannot promotes subunit exchange as it presumably cannot oligomerize with the B*.* subtilis GyrB domain of the fusion.

**Heterodimers mutated on two sides, such as BA_F_.A_59_; Contamination versus partial interface swapping and the role of GyrB flexibility and the free GyrB dimers.**

Heterodimer designed to have mutations on both sides have been purified by us and (Gubaev, Weidlich et al. 2016). Usually, the catalytic tyrosine is mutated on one side, which abolishes cleavage on this side. The other side bears another mutation, which abolishes supercoiling activity when present on both sides in a homodimer. The CTD is involved in wrapping the DNA in a positive loop prior to strand passage. Therefore, its complete absence from the gyrase complex will abolish supercoiling. However, losing the CTD on one side only of a dimer with two catalytic tyrosine does not abolish supercoiling activity as the dimer can use the single CTD for wrapping, as predicted by the strand-passage model. The supercoiling activity goes down only slightly, by a factor of two. However, if a catalytic tyrosine is mutated on the other side of the missing CTD, a drastic drop of supercoiling activity is observed, suggesting having two catalytic tyrosines is indeed important (see main discussion). These results were obtained in the following configuration: BA_59_.A_F_ + free GyrB. Similarly, one can mutate the ATP hydrolysis activity on one side (with the E44Q mutation on the GyrB subunit) and only slightly diminish the supercoiling activity, confirming earlier observations that hydrolysis of only one ATP is necessary for strand passage by type II topoisomerase (ref). The authors analyzed the analogous configuration: BA_Q_.A_F_ + free GyrB. Interestingly, the supercoiling activity observed is much less reduced that in the BA_59_.A_F_ + free GyrB configuration (we surmise that it would be still lower than the configuration with two catalytic tyrosine, although no quantitative measurements are provided). The swiveling model does not account for this difference, whereas interface swapping (IS) does since we have shown that having two CTDs is important for IS. The BA_Q_.A_F_ has both CTDs and therefore is expected to be more efficient for IS and consistently, shows more efficient reconstitution of supercoiling activity. The authors have also tested the configuration: BA_Q_.BA_F_, where both sides are BA fusion and no free GyrB is added. The supercoiling activity is much reduced, consistent with our observation that constraining GyrB flexibility diminishes supercoiling efficiently. In addition, the absence of free GyrB could also affect IS since we have shown that an excess of GyrB favors IS, potentially through oligomerization of the free GyrB subunits with the heterodimer constructs. Therefore, in the case of BA_Q_.BA_F_, the lack of free GyrB subunits could reduce efficiency of IS, thereby reducing the efficiency of supercoiling. Therefore, variation of IS amongst heterodimer constructs can underly their difference in supercoiling activity.

However, when these heterodimers undergo *complete* (we emphasize) subunit exchange, meaning all three interfaces are broken and exchange to reconstitute two gyrase dimer separated in solution, the resulting dimers are expected to be both inactive. This was argued to exclude complete subunit exchange as an explanation for the reconstitution of supercoiling by heterodimers with one catalytic tyrosine mutated. However, our experiments suggest that the DNA interface only is exchanged and probably occurs within multimers of the heterodimer, without producing free exchanging gyrase dimers. Therefore, in the case of BA_Q_.A_F_ and BA_Q_.BA_F_, it is possible that only the DNA gate is exchanged, whereas the GyrB subunits are not and retain strand capture activity within the oligomer. The oligomerization of a number of active gyrases could therefore result in a super-complex within which DNA gates can be exchanged, with strand-capture activity provided by a neighboring gyrase. It would be interesting to test our LLL mutant with the E44Q mutation introduced on the other side of the heterodimer. We would expect that the LLL mutation favors DNA gate interface swapping and therefore increase reconstitution of double-strand DNA cleavage. If this double-strand cleavage activity correlates with the reconstitution of supercoiling activity, we would conclude that the exchanged interface can use the ATPase from a neighboring complex for strand capture activity. In addition, interface swapping of a contaminating heterodimer including endogenous GyrA (like BA_Q_.A_endogenous_), expected to be efficient, could also account for the significant supercoiling activity observed. In contrast, in the case of the CTD mutant, which is on GyrA and away from the interface, the cleavage activity is reconstituted by exchanged DNA interfaces which are lacking CTDs on both side. Our results showed that this exchange does not result in increased supercoiling activity. Therefore, the exchanged DNA gates cannot use CTD that are not directly attached to the exchanged interface. We therefore conclude that, in the case of the CTD mutant, only interface swapping of a contaminating heterodimer including endogenous GyrA (BA_59_.A_endogenous_ or BA_F_.A_endogenoous_) accounts for the observed weak supercoiling activity. Again, the level of supercoiling roughly seems to correlate with the level of expected IS; although this would warrant careful quantitation and analysis.

Gubaev, A., D. Weidlich and D. Klostermeier (2016). "DNA gyrase with a single catalytic tyrosine can catalyze DNA supercoiling by a nicking-closing mechanism." Nucleic Acids Res **44**(21): 10354-10366.
